# A mutation in the *Drosophila melanogaster eve* stripe 2 minimal enhancer is buffered by flanking sequences

**DOI:** 10.1101/2020.06.19.162164

**Authors:** Francheska Lopez-Rivera, Olivia K. Foster, Ben J. Vincent, Edward C. G. Pym, Meghan D. J. Bragdon, Javier Estrada, Angela H. DePace, Zeba Wunderlich

**Affiliations:** Department of Systems Biology, Harvard Medical School, Boston, MA 02115; GSAS Research Scholar Initiative, Harvard University, Cambridge, MA 02138

**Keywords:** Drosophila melanogaster, enhancer, even-skipped, transcription factor binding site

## Abstract

Enhancers are DNA sequences composed of transcription factor binding sites that drive complex patterns of gene expression in space and time. Until recently, studying enhancers in their genomic context was technically challenging. Therefore, minimal enhancers, the shortest pieces of DNA that can drive an expression pattern that resembles a gene’s endogenous pattern, are often used to study features of enhancer function. However, evidence suggests that some enhancers require sequences outside the minimal enhancer to maintain function under environmental perturbations. We hypothesized that these additional sequences also prevent misexpression caused by a transcription factor binding site mutation within a minimal enhancer. Using the *Drosophila melanogaster even-skipped* stripe 2 enhancer as a case study, we tested the effect of a Giant binding site mutation (gt-2) on the expression patterns driven by minimal and extended enhancer reporter constructs. We found that, in contrast to the misexpression caused by the gt-2 binding site mutation in the minimal enhancer, the same gt-2 binding site mutation in the extended enhancer did not have an effect on expression. The buffering of expression levels, but not expression pattern, is partially explained by an additional Giant binding site outside the minimal enhancer. Mutating the gt-2 binding site in the endogenous locus had no significant effect on stripe 2 expression. Our results indicate that rules derived from mutating enhancer reporter constructs may not represent what occurs in the endogenous context.

## Introduction

Many developmental genes are expressed in complex patterns in space and time. The instructions for these patterns are largely encoded in enhancers, stretches of DNA composed of transcription factor (TF) binding sites. The earliest studies of enhancer function established that enhancers can retain their activity in synthetic reporter constructs, giving rise to the widely-held notion that enhancers are modules with distinct boundaries (Shlyueva *et al.* 2014). The idea that enhancers have distinct boundaries is reinforced by the way enhancers were traditionally identified – by reducing the DNA upstream of a gene’s promoter into increasingly small fragments until a “minimal” enhancer that was sufficient to produce all or a subset of a gene’s expression pattern was identified. Even when using modern functional genomic methods, enhancers are annotated with finite boundaries and attempts are often made to identify the minimal enhancer (Arnold *et al.* 2013; Koenecke *et al.* 2016; Diao *et al.* 2017; Monti *et al.* 2017).

Minimal enhancer reporter constructs have been a powerful tool for studying transcriptional control. By mutating minimal enhancers in reporters, scientists have identified key roles for transcription factor (TF) binding sites (Ney *et al.* 1990; Arnosti *et al.* 1996; Ma *et al.* 2000; Milewski *et al.* 2004). With the advent of high-throughput DNA synthesis and sequencing, this approach has been extended to study the effects of large numbers of enhancer variants in massively parallel reporter assays (Patwardhan *et al.* 2009; Melnikov *et al.* 2012; Inoue and Ahituv 2015; White 2015). An important, but often unstated assumption of this approach is: to decipher regulatory genetic variation in the intact genome, we can extrapolate from the measurements of variation in reporters driven by minimal enhancers, if we assume that enhancers are modular; in other words mutations would behave identically in an isolated enhancer and in the genome. Here, we set out to test this assumption directly.

There are several observations that enhancer function, particularly as defined by a minimal enhancer, may not be completely modular (Spitz and Furlong 2012; Lim *et al.* 2018). When measured quantitatively, the expression driven by some enhancer reporters does not precisely match the endogenous pattern (Staller *et al.* 2015). In many loci, the paradigm of a single enhancer driving expression in a single tissue is often an oversimplification. For example, in some loci, minimal enhancers cannot be identified for a given expression pattern, and many genes are controlled by seemingly redundant shadow enhancers (Barolo 2012; Sabarís *et al.* 2019). Furthermore, enhancer boundaries defined by DNAse accessibility and histone marks often do not match minimal enhancer boundaries defined by activity in reporters (Kwasnieski *et al.* 2014; Henriques *et al.* 2018). In some cases, the minimal enhancer is sufficient for an animal’s viability under ideal conditions, but sequences outside of the minimal enhancer are required for viability when the animal is exposed to temperature perturbations (Ludwig *et al.* 2011). Together, these examples highlight that while minimal enhancer regions can approximate the expression patterns of a gene, sometimes very closely, quantitative measurements of these regions’ activities can reveal their inability to recapitulate the nuances of gene regulation in the endogenous context.

In this work, we directly test the assumption that the misxpression caused by a mutation in a minimal enhancer reporter construct will also be observed when the same mutation is found in the genome. We compared the changes in gene expression caused by a mutation in three versions of an enhancer: 1) a minimal enhancer in a reporter, 2) an extended enhancer that contains the minimal enhancer plus flanking sequences in a reporter, and 3) in the endogenous locus. If the minimal enhancer truly represents a modular functional enhancer unit, the effects of the mutation on gene expression will be the same in each of these contexts. If not, the effects caused by the mutation will differ.

We use the well-studied *Drosophila melanogaster even-skipped (eve)* stripe 2 enhancer as our case study for several reasons (Goto *et al.* 1989; Small *et al.* 1992). *Eve* encodes a homeodomain transcription factor essential for proper segment formation in *Drosophila*, and five well-characterized enhancers drive its seven-stripe expression pattern in the blastoderm embryo (Figure 1A). To understand the mechanism of *eve* stripe 2 enhancer function, classic experiments mutated transcription factor binding sites in minimal enhancer reporter constructs, resulting in a set of variants with known effects that we can test in an extended enhancer construct and in the endogenous locus (Small *et al.* 1992; Arnosti *et al.* 1996). Subsequent experiments showed that, while the *eve* stripe 2 minimal enhancer is sufficient for an animal’s viability in *D. melanogaster*, the sequences outside of minimal enhancer are required to drive robust patterns of gene expression when the animal is exposed to temperature perturbations (Ludwig *et al.* 2011), or to drive a proper stripe in other species (Crocker and Stern 2017). Together, these experiments indicate that the minimal enhancer does not recapitulate the complete transcriptional control of *eve* stripe 2. The *Drosophila* blastoderm embryo also provides technical advantages; we can readily incorporate reporter constructs, make genomic mutations, and measure levels and patterns of gene expression at cellular resolution (Hendriks *et al.* 2006; Wunderlich *et al.* 2014). This allows us to measure potentially subtle differences in expression patterns and levels driven by different enhancer variants.

**Figure 1:**
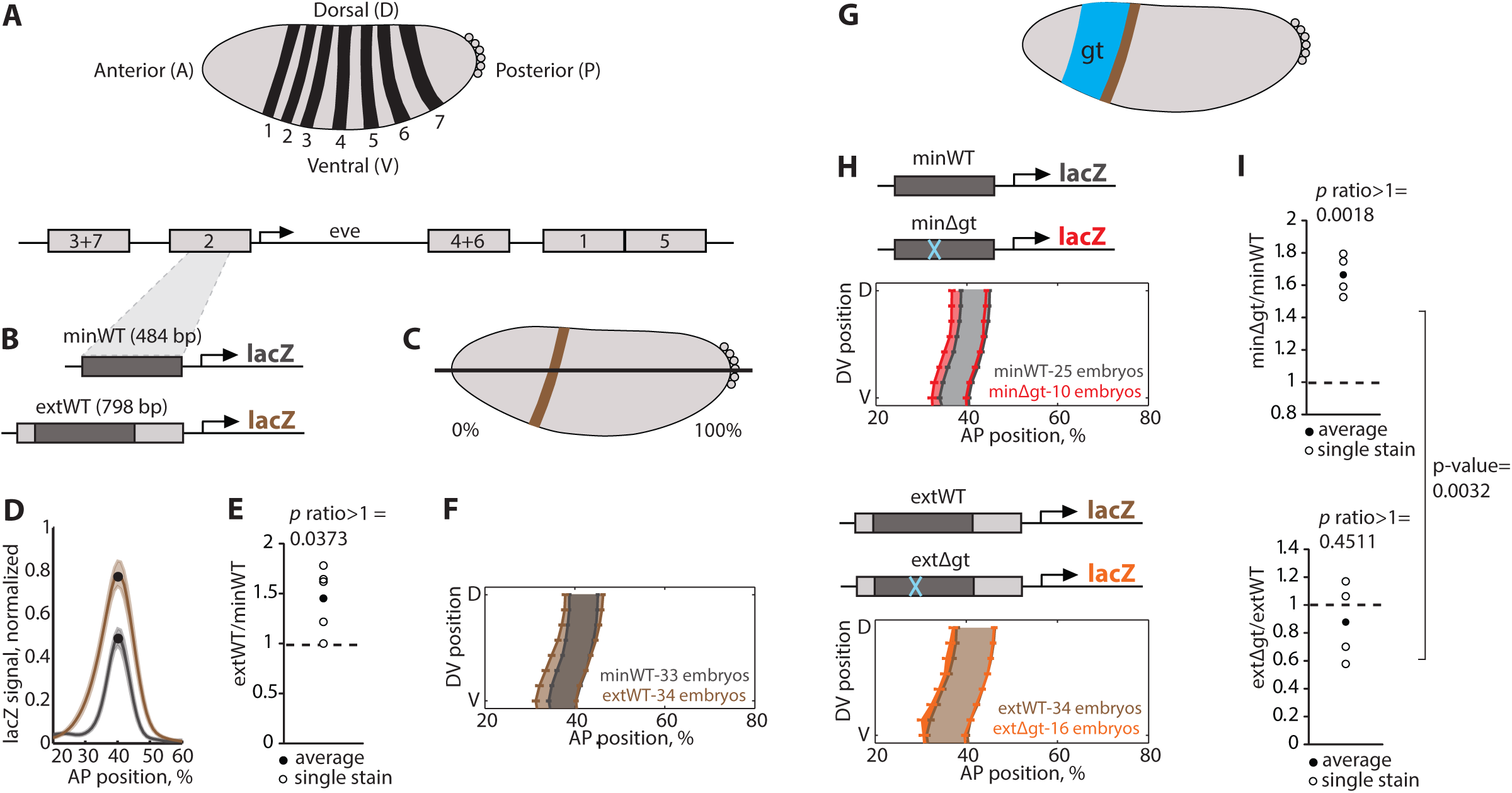
The effect of the *gt-2* binding site mutation is buffered in the *eve* stripe 2 extended enhancer. (A) *Eve* is expressed as a pattern of seven stripes along the anterior-posterior axis of the *Drosophila melanogaster* blastoderm, and this pattern is driven by five enhancers. (B) We generated transgenic reporter fly lines with the wild-type minimal (minWT) and extended (extWT) *eve* stripe 2 enhancers, and we measured *lacZ* expression in embryos using *in situ* hybridization. (C, D) We plotted *lacZ* levels in a lateral strip of cells along the AP axis for the minWT (dark gray) and extWT (brown) enhancers measured in a single stain, with the shading showing the standard error of the mean. The extWT enhancer drives a higher peak level of expression. (E) We calculated the ratio of peak *lacZ* expression levels (black dots in D) driven by the extWT and minWT enhancers in five different stains (open circles). The average ratio of the five stains is represented by a closed circle. The extWT enhancer drives 1.45 times higher expression than the minWT (p-value extWT/minWT > 1 = 0.0373). (F) We show the average boundary positions of the *lacZ* expression pattern. Error bars show standard error of the mean boundary positions of the expression pattern. The extWT enhancer drives a wider pattern of expression (brown shading) than the minWT enhancer (gray shading), with the anterior border of the stripe laying ∼1.6 cell widths more anterior than the minWT enhancer pattern. (G) The transcription factor Giant (Gt) is expressed as a broad band anterior to *eve* stripe 2 and represses *eve*, establishing the anterior boundary of stripe 2. (H, I) We characterized the expression patterns and levels driven by the minimal (minΔgt, top panels) and extended (extΔgt, bottom panels) enhancers with a gt-2 binding site deletion. In the minΔgt enhancer, the gt-2 deletion causes an anterior shift in the anterior boundary of the expression pattern and an increase in expression level. In the extended enhancer, the gt-2 deletion causes a more modest shift in the anterior boundary and no significant change in peak expression level.

We hypothesized that a transcription factor binding site mutation will have its maximum effect on gene expression when found in a minimal enhancer, while its effects will be reduced, or buffered, when found in the extended enhancer and in the endogenous locus due to the contributions of additional regulatory DNA sequences. We tested our hypothesis and found that the effects of a TF binding site mutation on gene expression are indeed buffered in the extended *eve* stripe 2 enhancer and in the endogenous locus. This buffering is partially explained by an additional binding site in the sequence outside the *eve* stripe 2 minimal enhancer. These results imply that we cannot always extrapolate the effects of enhancer mutations in minimal reporters to extended sequences or to the endogenous intact locus. We discuss implications of our results for studying the functional consequences of regulatory sequence variation.

## Materials and Methods

### Enhancer sequences and mutations in reporter constructs

Each of the *eve* stripe 2 enhancer sequences was cloned into a pB*ϕ*Y plasmid containing an *eve* basal promoter-*lacZ* fusion gene, the *mini-white* marker, and an attB integration site. The enhancer sequences are located immediately upstream of the *eve* basal promoter. All constructs were integrated by Genetic Services, Inc. into the attP2 docking site of the *Drosophila melanogaster y*[1], *w*[67c23] line. We followed the *mini-white* eye marker as we conducted crosses to make the transgenic fly lines homozygous.

The 484 base pair (bp) wild-type minimal (minWT) enhancer sequence was defined by Small and colleagues (Small *et al.* 1992). MinΔgt is the minWT enhancer with a 43 bp deletion of the giant-2 (gt-2) binding site as described in (Small *et al.* 1992). The wild-type extended (extWT) enhancer is the minWT sequence plus the 50 bp upstream and 264 bp downstream flanking sequences present in the *eve* locus. The boundaries of the extWT enhancer are two conserved blocks of 18 and 26 bp on the 3’ and 5’ ends of the enhancer (Ludwig *et al.* 1998). The extΔgt enhancer consists of the extWT enhancer with the same gt-2 binding site deletion as in minΔgt.

To computationally predict additional Gt sites in the extended enhancer, we used PATSER and three different Gt position weight matrices (PWMs) generated with data from yeast one-hybrid, DNA footprinting, and SELEX assays (Hertz and Stormo 1999; Noyes *et al.* 2008; Li *et al.* 2011; Schroeder *et al.* 2011). A common Gt binding site was found in the downstream flanking sequence of the extended enhancer using all three PWMs with a p-value of 0.001. Because of overlaps with other predicted binding sites, this Gt binding site was mutated by changing five nucleotides in extΔgt to create the extΔgt,Δgt enhancer.

The minWT-sp1 and minWT-sp2 enhancers consist of the minWT enhancer and two different 264 bp downstream spacer sequences, sp1 and sp2. Each of these sequences are about half of a 500 bp lacZ sequence from which we removed high affinity binding sites for Bicoid, Hunchback, Giant, and Kruppel, using a PATSER p-value of 0.003. The minΔgt-sp1 enhancer is composed of minΔgt and sp1. MinΔgt-sp1+gt is the minΔgt-sp1 enhancer containing the additional *gt* binding site that we identified, located in the position where it is found in the extended enhancer. File S1 contains the sequences of all the enhancers that were tested in reporter constructs.

### Endogenous eve giant-2 deletion using the CRISPR system

Briefly, gRNAs (5’-TCTAACTCGAAAGTGAAACGAGG-3’ and 5’-ATTCCGTCTAAATGAAAGTATGG-3’) adjacent to the gt-2 binding site were cloned into pU6-BbsI-chiRNA. A ScarlessDsRed selection cassette (https://flycrispr.org/scarless-gene-editing/) was used with ∼500 bp homology arms flanking the gRNA cut sites in the *eve* stripe 2 enhancer. These plasmids were injected into *y*[1] *w*[67c23]; attP2{nos-Cas9} by BestGene. The dsRed selection cassette was mobilised by crossing to *w*[1118]; In(2LR)Gla, *wg*[Gla-1]/CyO; Herm{3xP3-ECFP, alphatub-piggyBacK10}M10, and selecting for non-dsRed eyed flies, to give the final allele *eve*[ahd4]. Further crosses to remove the transposase yielded flies with the genotype *w*[1118]; *eve*[ahd4], which we term “Δgt *eve* locus.” The edit was confirmed by PCR. The control flies to which the CRISPR flies were compared had the genotype *y*[1] *w*[67c23]; attP2{hbP2-LacZ}.

### In situ hybridization and imaging

We collected and fixed 0-4 hour old embryos grown at 25°C, and we stained them using *in situ* hybridization as in (Hendriks *et al.* 2006; Wunderlich *et al.* 2014). We incubated the embryos at 56°C for two days with DNP-labelled probes for *hkb* and DIG-labelled probes for *ftz*. Transgenic reporter embryos were also incubated with a DNP-labeled probe for *lacZ*, and the WT *eve* locus and Δgt *eve* locus CRISPR embryos were incubated with a DNP-labeled probe for *eve. Hkb* probes were used to normalize *lacZ* expression levels between the different transgenic reporter lines. The DIG probes were detected with anti-DIG-HRP antibody (Roche, Indianapolis, IN) and a coumarin-tyramide color reaction (Perkin-Elmer, Waltham, MA), and the DNP probes were detected afterwards with anti-DNP-HRP (Perkin-Elmer) antibody and a Cy3-tyramide color reaction (Perkin-Elmer). Embryos were treated with RNAse and nuclei were stained with Sytox green. We mounted the embryos in DePex (Electron Microscopy Sciences, Hatfield, PA), using a bridge of #1 slide coverslips to avoid embryo morphology disruption.

Reporter embryos from the early blastoderm stage (4-10% membrane invagination, roughly 10-20 minutes after the start of the blastoderm stage) were imaged, and CRISPR embryos from early blastoderm stage (9-15% membrane invagination, roughly 15-25 min after the start of the blastoderm stage) were imaged. We used 2-photon laser scanning microscopy to obtain z-stacks of each embryo on a LSM 710 with a plan-apochromat 20X 0.8 NA objective. Each stack was converted into a PointCloud, a text file that includes the location and levels of gene expression for each nucleus (Hendriks *et al.* 2006).

### Data analysis of reporter constructs

To normalize the *lacZ* levels in the reporter embryos, we divided the *lacZ* signal by the 95% quantile of *hkb* expression in the posterior 10% of each embryo (Wunderlich *et al.* 2014). We expect the *lacZ* and *hkb* levels to be correlated within a transgenic line. To verify this, we ran a regression of the 99% quantile *lacZ* value from each embryo and the 95% quantile *hkb* value. Cook’s distance (Cook, 1977) was used to discard influential outliers (Wunderlich *et al.* 2014). To avoid extraneous sources of noise in the normalization, we only compare *lacZ* levels between embryos with the same genetic background and stained in the same *in situ* hybridization experiment.

To calculate the average *lacZ* expression levels along the anterior-posterior (AP) axis in each transgenic line, we used the extractpattern command in the PointCloud toolbox, which can be found in http://bdtnp.lbl.gov/Fly Net/bioimaging.jsp?w=analysis. This command divides the embryo into 16 strips around the dorso-ventral (DV) axis of the embryo, and for each strip, calculates the mean expression level in 100 bins along the anterior-posterior (AP) axis. We averaged the strips along the right and left lateral sides of the embryos and subtracted the minimum value along the axis to remove background noise.

We calculated the peak average *lacZ* expression level within the *eve* stripe 2 region for each transgenic line in each *in situ* experiment separately. We then calculated the ratio between the peak average *lacZ* expression levels of two transgenic lines stained in the same *in situ* experiment. Ratios were calculated for each stain and the average ratio from multiple stains was determined. Comparisons between average ratios and 1 or between two different ratios were made by using one- or two-sample t-tests with unequal variances.

The boundaries of *eve* stripe 2 expression were defined as the inflection point of the *lacZ* expression levels. Since the boundaries of lacZ expression should not change between stains, plots with the average boundaries of *lacZ* expression in each transgenic line were made with embryos pooled from multiple stains (see Figure S2 for number of embryos measured for each genotype). The cell length differences were calculated by determining the average position of the boundary across the DV axis of the embryos analyzed. One cell length is approximately equivalent to one percent of the embryo length.

### Data analysis of endogenous eve giant-2 deletion

Briefly, we normalized to *eve* stripe 1 cellular expression to compare *eve* levels in the *eve*[ahd4] embryos and the control (Fowlkes *et al.* 2008). As described above, using the extractpattern command from the PointCloud toolbox, we found an averaged lateral trace across both sides of the embryo. The peak average *eve* expression for each stripe was normalized to the peak average expression of *eve* stripe 1. We performed a comparison of stripe levels between conditions using a two-sided rank sum test.

The boundary of *eve* stripes were defined as above using extractpattern and, for a given embryo, eight boundary positions on the left and right lateral sides were averaged. Plots with the average boundary of *eve* stripe 2 in the *eve*[ahd4] versus control were made with embryos pooled from different stains. To compare boundaries between the two genotypes, a Mann-Whitney U Test was used, with the factors being one of the eight dorso-ventral positions along both lateral sides of the embryo and the embryo genotype. The p-value was reported for the genotype factor effect.

### Data Availability

All transgenic and CRISPR fly lines are available upon request. File S1 contains the sequences for all enhancer constructs. Figure S2 contains all ratios presented in Figures 1-3 in one plot. Figure S5 contains the enhancer sequence of the *eve*[ahd 4] locus as well as a map of the predicted binding sites.

**Figure 2:**
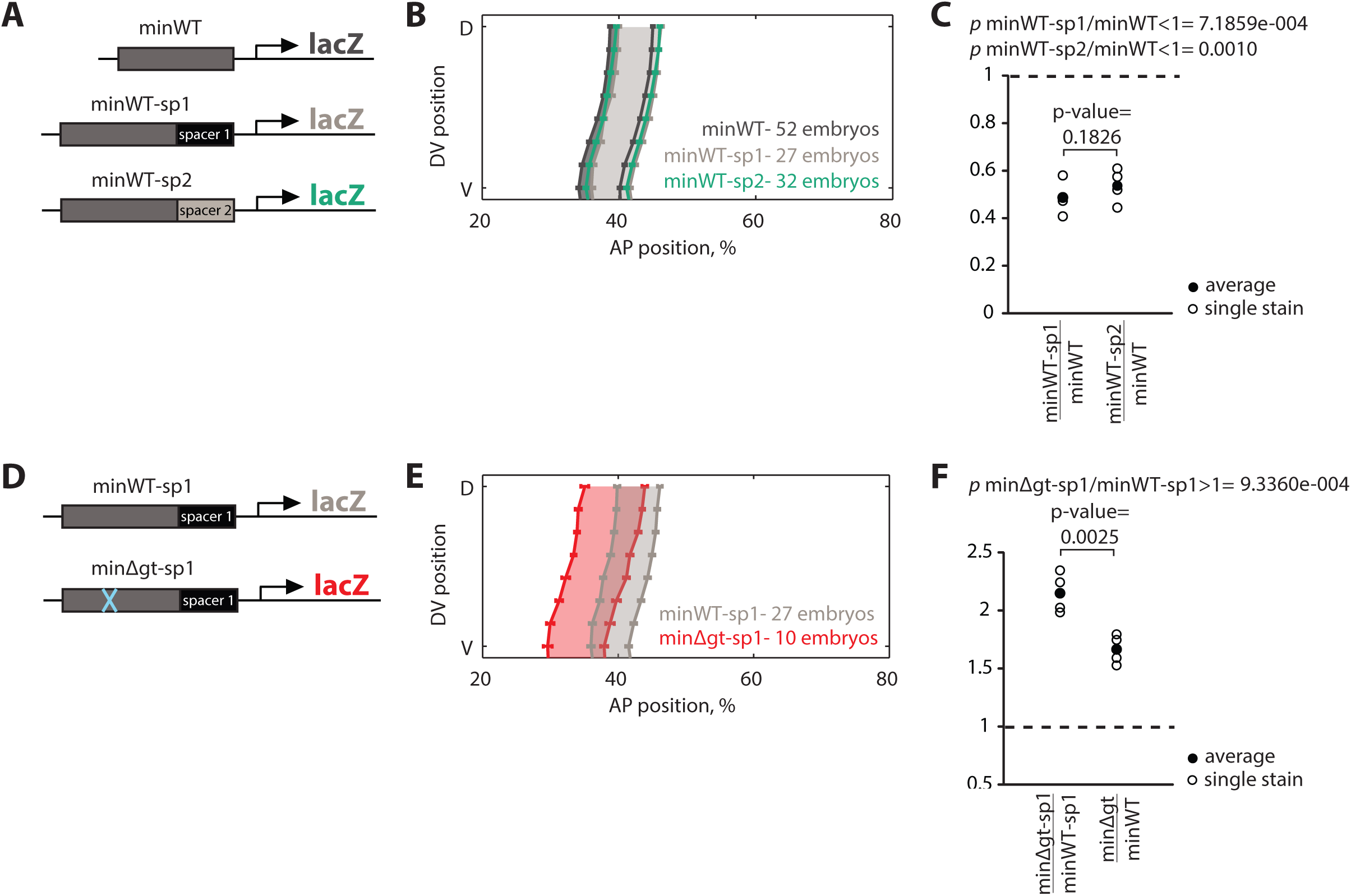
Distance from the promoter reduces *eve* stripe 2 expression levels and is not sufficient to explain the buffering. (A) To test if distance from the promoter contributes to buffering the gt-2 deletion, we used two different 264 bp spacer sequences (sp1 and sp2) to make two constructs, minWT-sp1 and minWT-sp2. (B) We find that moving the minimal enhancer away from the promoter slightly shifts the boundaries of the stripe to the posterior. Error bars show standard error of the mean boundary positions of the expression pattern. (C) A comparison of peak expression levels shows that moving the minimal enhancer away from the promoter reduces peak expression levels. (D) We tested if distance from the promoter is sufficient to explain the Gt site deletion buffering in the extended enhancer by introducing the Gt site deletion into the minWT-sp1 construct, minΔgt-sp1. (E) The minΔgt-sp1 construct drives an expression pattern that is dramatically shifted to the anterior, indicating that the spacer cannot buffer the Gt binding site deletion’s effect on expression pattern. (F) The minΔgt-sp1/minWT-sp1 peak expression ratio is significantly larger than minΔgt/minWT ratio, indicating that the gt-2 deletion has a more dramatic effect in the minΔgt-sp1 and that the increasing distance from the promoter does not buffer the effects of the gt-2 deletion.

**Figure 3:**
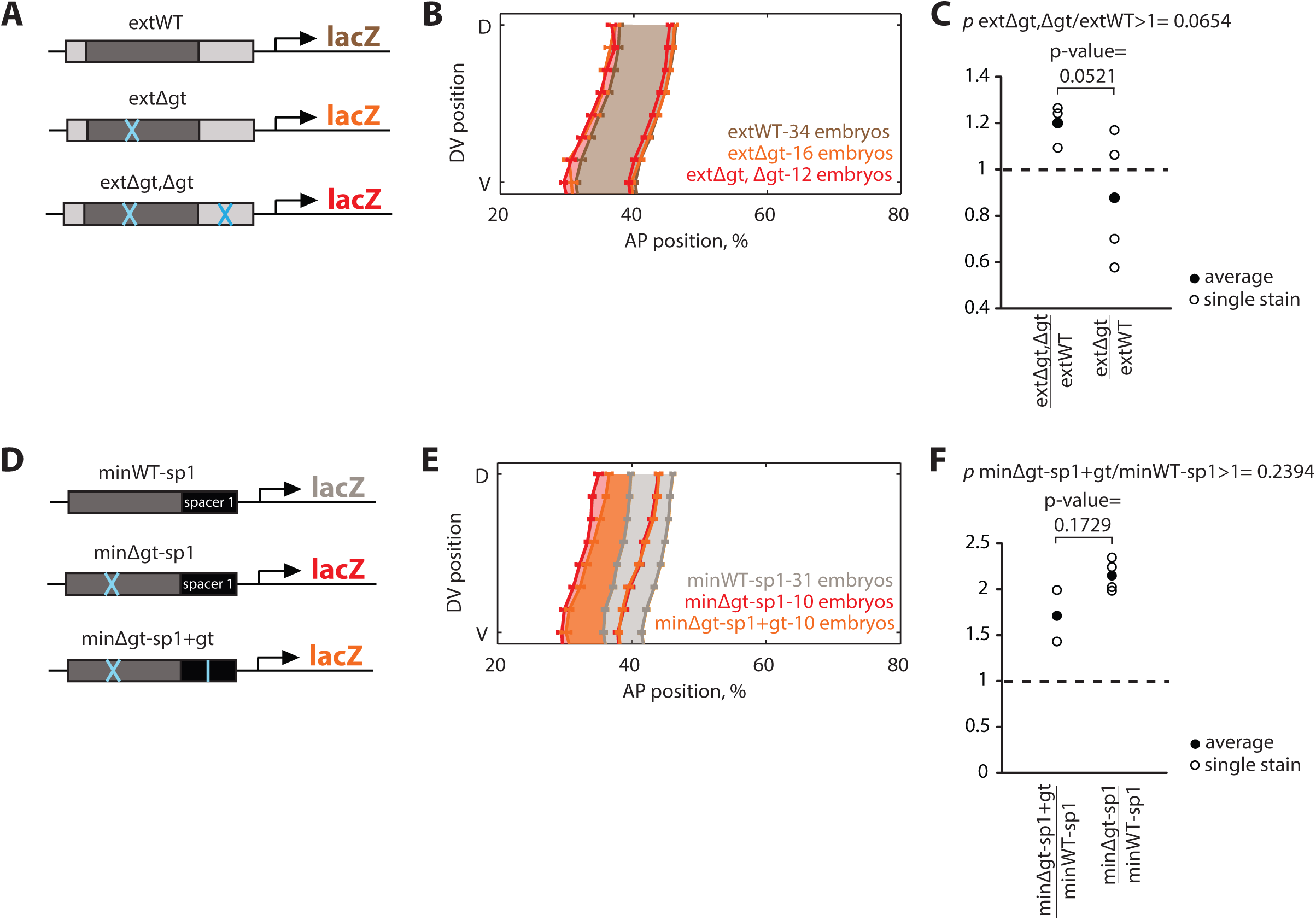
An additional Gt binding site partially explains the buffering. (A) We found an additional predicted Gt binding site outside the minimal enhancer sequence but within the extended enhancer. A reporter construct, extΔgt,Δgt, testing for the necessity of the additional Gt binding site was made by mutating the predicted Gt binding site. (B) The average position of the *lacZ* anterior boundaries was nearly identical for the extΔgt and extΔgt,Δgt constructs, indicating that eliminating the additional Gt binding site does not affect buffering of the gt-2 deletion of the expression pattern. Error bars show standard error of the mean boundary positions of the expression pattern. (C) If the additional *gt* site was necessary and sufficient for the buffering, the extΔgt,Δgt/extWT ratio would be higher than 1 and very similar to minΔgt/minWT ratio. If the additional Gt binding site was not necessary at all, the extΔgt,Δgt/extWT ratio would be similar to 1 and to the extΔgt/extWT ratio. The results suggest that the additional Gt site explains only some of the buffering of the gt-2 deletion in level. (D) We tested the sufficiency of the additional Gt binding site by making a construct with the minΔgt-sp1 element and inserting the additional Gt binding site in the same location as it is found in the extended enhancer (minΔgt-sp1+gt). (E) The additional Gt binding site shifts the anterior boundary of expression slightly to the posterior, when compared to the pattern driven by minΔgt-sp1. (F) The peak minΔgt-sp1+gt/minWT-sp1 ratio is lower, but not significantly different from the minΔgt-sp1/minWT-sp1 ratio, indicating that this Gt binding site is not sufficient to explain the buffering of expression level in the extended enhancer.

## Results

### The minimal and extended eve stripe 2 enhancers drive different patterns and levels of expression

To test the effects of mutations in the minimal and extended *eve* stripe 2 enhancer on expression, we began by characterizing the wild type (WT) expression patterns driven by the previously-defined minimal (minWT) and extended (extWT) enhancers (Figure 1B-F). The minimal enhancer is 484 bp and was identified as the smallest piece sufficient to drive expression in the region of stripe 2 (Small *et al.* 1992). The extended enhancer boundaries were chosen as the two conserved blocks of 18 and 26 bp on the 3’ and 5’ sides of the minimal enhancer, resulting in a 798 bp piece (Ludwig *et al.* 1998). We generated transgenic animals with *lacZ* reporter constructs inserted into the same location of the genome, and we measured *lacZ* expression using *in situ* hybridization and a co-stain for normalization (Wunderlich *et al.* 2014). The stripe driven by the extended enhancer is wider – its anterior boundary is ∼1.6 cell widths more anterior than that of the minimal enhancer (Figure 1F). In addition, the peak *lacZ* expression driven by the extWT is 1.45 times higher than the minWT enhancer (p-value ratio > 1 is 0.0373; Figure 1D, E).

### The gt-2 transcription factor binding site mutation is buffered in the extended enhancer

To test the effect of mutations in the minimal and extended enhancers, we looked to the literature to find a known sequence mutation that had a measurable effect on expression in the minimal enhancer. Previous work identified three footprinted binding sites within the minimal enhancer for the repressor Giant (Gt), which is expressed anterior of *eve* stripe 2 (Figure 1G). A minimal enhancer with a deletion of one of these binding sites, gt-2, drives higher and broader expression than the WT enhancer (Arnosti *et al.* 1996). We created reporters with the same deletion of gt-2 in the minimal and extended enhancers (Figure 1H) and measured the effect of the deletion on both expression levels and patterns. Consistent with previous results, we found that minΔgt drives 1.67 times the expression of the minWT enhancer (p-value expression ratio > 1 is 0.0018 in Figure I, top), and a pattern that is expanded 1.7 cell widths to the anterior (Figure H, top). In contrast, the expression level driven by the extΔgt enhancer is not significantly different from the extWT enhancer (p-value expression ratio > 1 is 0.4511 in Figure 1I, bottom), and the pattern is expanded by only 0.9 cell widths (Figure 1H, bottom). The minΔgt/minWT expression ratio is also significantly larger than the extΔgt/extWT ratio (p-value=0.0032), indicating that the deletion has a much larger effect on the expression level driven by the minimal enhancer than the extended enhancer. Together, these results indicate that the effect of the gt binding site deletion is buffered in the extended enhancer.

### Distance from the promoter reduces expression levels and does not explain buffering

The minimal and extended enhancers differ from one another in the flanking sequences. These flanks may contribute to buffering in two primary ways: 1) the flanks may contain TF binding sites or other specific sequence elements, and 2) the flanks increase the distance of the minimal piece from the promoter.

In the minWT constructs the enhancer is 38 bp from the promoter, whereas in the extWT constructs the same minWT sequence is located 302 bp away from the promoter. To test how this change in distance contributes to the differences in expression of the two constructs, we inserted two different 264 bp spacer sequences (sp1 and sp2) into the minWT reporters, to make the constructs minWT-sp1 and minWT-sp2 (Figure 2A). sp1 and sp2 are lacZ sequences from which high affinity binding sites for the regulators involved in eve stripe 2 expression have been removed. For both spacers, increasing the distance of the minWT sequence significantly reduces expression levels (0.48 for sp1, p = 7.189e-4, and 0.54 for sp2, p = 0.0010, Figure 2C), while only minimally affecting the AP positioning, with both showing marginal shifts to the posterior in comparison to minWT. The anterior and posterior boundaries of the minWT-sp1 are shifted to the posterior part of the embryo by 1.4 and 1.3 cell lengths, respectively, when compared to minWT (Figure 2B). The anterior and posterior boundaries of minWT-sp2 are shifted to the posterior by 1.0 and 1.1 cell lengths, respectively, when compared to minWT (Figure 2B). These data demonstrate that the level of expression driven by minWT is influenced by enhancer-promoter distance.

To test if promoter-enhancer distance explains the buffering of the gt*-*2 deletion, we made a construct with the minΔgt enhancer separated from the promoter by sp1, minΔgt-sp1, and compared it to minWT-sp1 (Figure 2D). If the distance from the promoter contributes to the buffering effect, the expression ratio of minΔgt-sp1/minWT-sp1 would be smaller than that of minΔgt/minWT, and the spatial pattern between minWT-sp1 and minΔgt-sp1 would be more similar than between minWT and minΔgt. In fact, the opposite is true – the ratio is larger (p-value=0.0025) and the spatial pattern is less similar, indicating that the relative distance of the core 484 bp to the promoter does not contribute to the buffering in the extended piece (Figure 2E, F).

### An additional Gt binding site in the flanking sequence partially explains the buffering

Since promoter-enhancer distance does not explain the buffering of the extended enhancer, the buffering must be due to differences in the sequence content of the minimal and extended enhancers. We hypothesized that there might be additional gt binding sites in the flanks of the extended enhancer that explain the observed buffering of the gt-2 deletion. We scanned these flanking regions with several existing Gt position weight matrices (PWMs) and found one binding site common to all the PWMs (see Materials and Methods and Figure S1). We mutated the common site to make the extΔgt,Δgt construct (Figure 3A). If this common site is responsible for the buffering, we would expect that the extΔgt,Δgt construct would drive higher expression levels and a wider stripe than the extWT construct. The extΔgt,Δgt enhancer drives a pattern with an anterior boundary that is not significantly different from the extΔgt enhancer (Figure 3B). Compared to the peak expression levels driven by the extWT enhancer, the extΔgt,Δgt enhancer drives 1.20 times the expression (p-value that ratio >1 is 0.0654) (Figure 3C). Because the peak expression ratio of extΔgt,Δgt/extWT is between that of minΔgt/minWT and extΔgt/extWT, this result suggests that the additional Gt binding site is partially responsible for buffering the effect of the gt-2 mutation on expression levels (Figure S2). However, since the extΔgt and extΔgt,Δgt enhancers drive virtually the same expression pattern, this binding site is not responsible for buffering the effect of gt-2 deletion on expression pattern. Therefore, this additional Gt binding site can only partially explain why the extended enhancer can buffer the effect of the gt-2 deletion. Additional Gt binding sites, other TF binding sites, or other functional sequences in the extended enhancer sequence flanks may be responsible for the unexplained buffering (see Discussion).

### Adding a Gt binding site to the minimal enhancer is not sufficient to buffer a Gt mutation

Since the additional Gt site is necessary to partially buffer the gt-2 deletion, we wanted to test whether it was also sufficient. We inserted the additional Gt binding site into the spacer of the minΔgt-sp1 construct in the same position as it is found in the extWT construct to make the minΔgt-sp1+gt construct (Figure 3D). We compared its expression to the minWT-sp1 and the minΔgt-sp1 constructs. If the additional Gt site is sufficient to buffer the gt-2 deletion, we would expect that the minΔgt-sp1+gt would drive lower expression levels than minΔgt-sp1 and a similar expression pattern to the minWT-sp1 construct. We found that the peak expression ratio of minΔgt-sp1+gt/minWT-sp1 was on average lower, but not significantly different from the minΔgt-sp1/minWT-sp1 ratio, indicating that this binding site alone is not sufficient to buffer the gt-2 deletion (p-value=0.1729) (Figure 3F). The expression patterns driven by minΔgt-sp1+gt and minΔgt-sp1 are also very similar, though there is a slight posterior shift of the anterior boundary in the minΔgt-sp1+gt construct (Figure 3E). It is possible that this binding site needs its original context to function properly, which may be due to the importance of binding site flanks on DNA shape (Rohs *et al.* 2010; Li and Eisen 2018), or other, unknown requirements.

### The gt-2 transcription factor binding site mutation is buffered in the endogenous locus

To test whether the gt-2 deletion can be buffered in the intact locus, as it is in the extended enhancer, we used CRISPR editing to generate flies homozygous for the same gt-2 deletion in the endogenous *eve* locus, which we called Δgt *eve* locus (Figure 4A, Figure S5). We then measured *eve* expression patterns and levels using *in situ* hybridization in the Δgt *eve* locus embryos and WT *eve* locus embryos (see Methods for details). To measure expression levels in *eve* stripe 2, we internally normalized to the levels of *eve* stripe 1, which is the first *eve* stripe to be expressed in this developmental stage (Figure S3; see Methods for details). We observed that the expression levels of *eve* stripe 2 in embryos with Δgt *eve* locus are not significantly different from those in embryos with WT *eve* locus (p-value=0.1007) (Figure 4C). The *eve* stripe 2 patterns driven by the WT *eve* locus and the Δgt *eve* locus are not significantly different (Mann-Whitney U test, Figure 4B). This suggests that the gt-2 deletion in the endogenous *eve* stripe 2 enhancer is buffered: expression levels and boundary position in the Δgt *eve* locus embryos are not significantly different from the WT *eve* locus embryos, in agreement with the observations made in the extended enhancer. Interestingly, we observed differences between Δgt *eve* locus and WT *eve* locus on other stripes of the *eve* pattern (Figure S4). There are differences in the expression levels of *eve* stripes 5 and 6, and in the patterns of *eve* stripe 4. We speculate that the differences might be due to the effects of the genetic backgrounds of Δgt *eve* and WT *eve* locus embryos (see Methods). All together, these results suggest that the effect of a specific mutation in the *eve* stripe 2 minimal reporter construct is not recapitulated when tested in the endogenous enhancer context.

**Figure 4:**
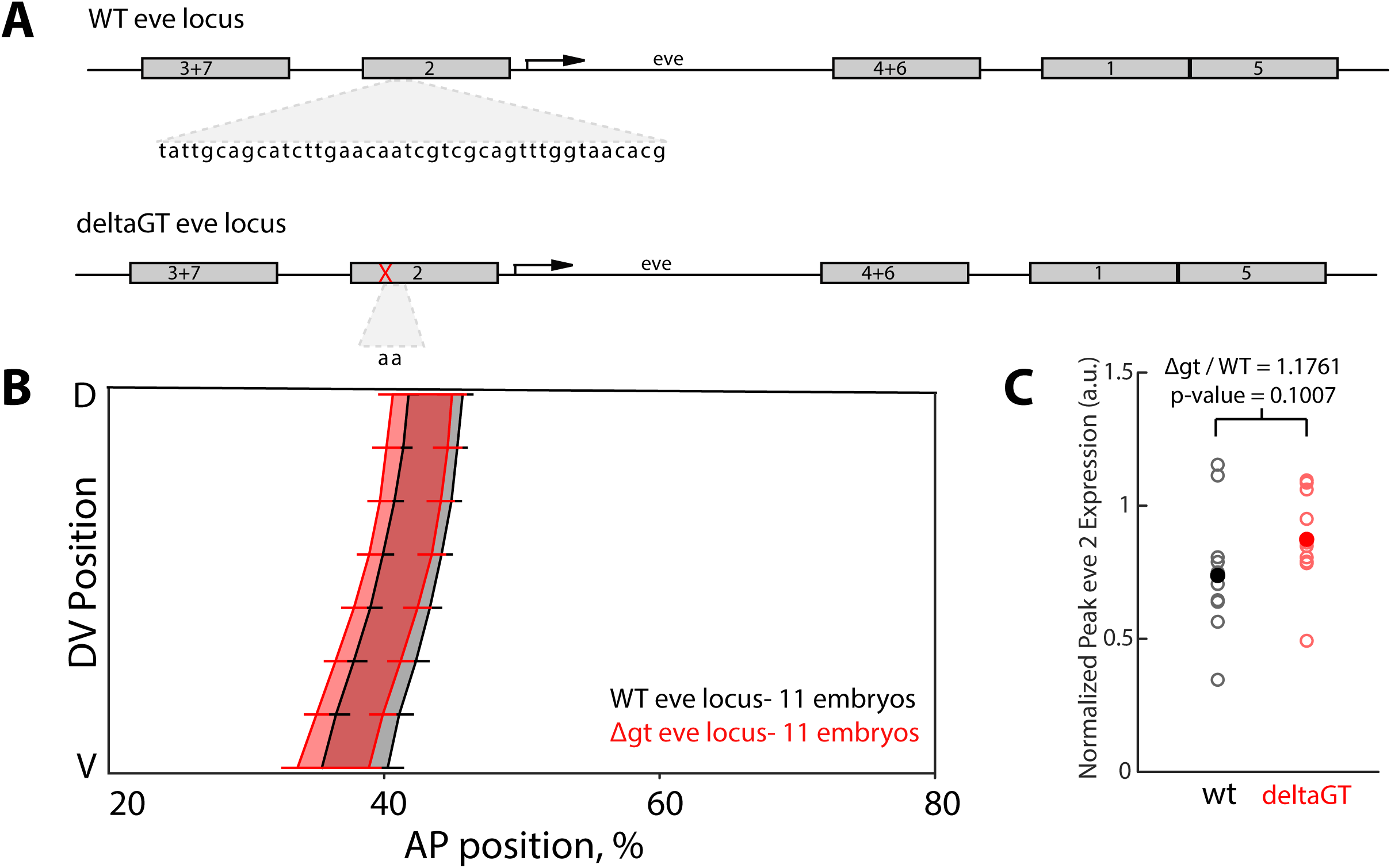
CRISPR deletion of Gt binding site in endogenous *eve* locus does not change stripe 2 expression. (A) Using a scarless-CRISPR method, we removed the gt-2 binding site endogenously. (B) The boundaries of *eve* stripe 2 are not significantly different between the Δgt *eve* locus embryos and the WT embryos (Mann-Whitney U test). Error bars show standard error of the mean boundary positions of the expression pattern. This indicates that the boundary of *eve* stripe 2 in the endogenous context is buffered against the removal of gt-2. (C) Normalized peak expression levels of *eve* stripe 2 did not change significantly in the Δgt *eve* locus versus control (p=0.1007). Moreover, the ratio of Δgt *eve* locus to WT *eve* locus equals 1.1761. Filled circles represent mean expression level and open circles are *eve* peak expression for each individual embryo analyzed (Δgt locus: n=11, control locus: n=11).

## Discussion

The desire to define discrete minimal sequences that are sufficient to drive gene expression patterns emerged from a combination of the technical limitations imposed upon early studies and the resulting “founder fallacy” (Halfon 2019), cementing the first discovered examples of enhancers into generalisations. Understanding and acknowledging the ways in which the activity of minimal enhancers in reporter constructs differs from the activity of the same sequences within the endogenous locus will help us understand gene regulatory logic at a genome scale, as well as regulatory variation and evolution; simultaneously, it reaffirms the important contributions that reporter constructs can still make to deciphering the mechanisms of transcription.

Using one of the textbook examples of an enhancer, *eve* stripe 2, we have shown that deletion of a key TF binding site for Gt has significant functional effects on the expression driven by the minimal enhancer sequence, but not when this minimal enhancer is modestly extended, nor when the same binding site is removed from the endogenous locus. Furthermore, we identified an additional Gt binding site found outside the minimal enhancer that contributes to buffering the effect of this mutation. However, this additional site partially buffers changes in expression level, but not position, and is necessary, but not sufficient to explain the buffering effect observed in the extended enhancer.

Given that there were no previously described Gt binding sites in the region flanking the minimal enhancer, it was somewhat unexpected that the effect of the gt-2 deletion would be buffered in the extended enhancer (Ludwig *et al.* 2011). However, finding all transcription factor binding sites remains a challenge and may explain why we cannot fully account for the gt-2 deletion buffering in the extended enhancer (Keilwagen *et al.* 2019). Gt’s binding preference has been measured using several techniques, which all yield different sequence motifs (Noyes *et al.* 2008; Li *et al.* 2011; Schroeder *et al.* 2011). We searched for Gt binding sites with three different sequence motifs, and we found and mutated a single high-affinity binding site predicted by all three motifs. But, there are additional predicted Gt binding sites that may be contributing to the buffering (Figure S1).

We do not understand why the minimal spacer constructs that include the gt-2 deletion show a large anterior shift of the posterior boundary of the expression pattern. The shift is not observed in the minimal spacer constructs that exclude the deletion or in the minΔgt or extΔgt constructs, so it is not due to the spacer sequence or to the gt-2 mutation individually. We hypothesize there is a specific promoter-enhancer interaction that occurs when both the spacer and the gt-2 deletion are present, but we cannot speculate on the precise underlying cause of this interaction.

This simple case study illustrates clearly that the effects of mutations, as measured in minimal enhancer sequences, cannot be simply extrapolated to larger enhancer regions or to the enhancer in its endogenous context in the genome. These results provide additional evidence challenging the idea that enhancers are strictly modular and that they have defined boundaries (Evans *et al.* 2012; Halfon 2019; Sabarís *et al.* 2019). Experiments using minimal enhancer reporter constructs have been extremely valuable for identifying genetic interactions and mechanisms of transcriptional control, e.g. activator/repressor balance and short- and long-range repression (Arnosti *et al.* 1996; Kulkarni and Arnosti 2005; Vincent *et al.* 2018). However, as more high throughput methods are developed to test the effect of mutations in small to medium-size enhancer fragments, we need to be cautious in interpreting these results (Inoue and Ahituv 2015). A mutation that may have dramatic effects on expression when made in a minimal enhancer may have no effect when made in the genome of an animal.

To test the mutation effects definitively, reporter construct experiments need to be complemented with manipulations of the endogenous enhancer sequences. Due to the CRISPR revolution, these types of experiments are becoming increasingly feasible (Zhou *et al.* 2014; Kvon *et al.* 2016; Rogers *et al.* 2017), and methods are being developed to use high-throughput CRISPR experiments to identify and perturb enhancers, as reviewed in (Lopes *et al.* 2016; Catarino and Stark 2018). These experiments will provide the data to attack the challenge of modeling the function of increasingly large pieces of the genome simultaneously, which is ultimately required to predict how variation in enhancer sequences affects gene expression.

## Supporting information

Compiled Supplementary Material

## Acknowledgements

We thank all the members of the DePace lab for helpful discussions and suggestions on the manuscript. The research reported in this publication was funded by the Harvard GSAS Research Scholar Initiative (to F.L.R.), NIH grants K99/R00 HD073191 and R01 HD095246 (to Z.W.), the NSF Graduate Research Fellowship DGE1745303 (to O.K.F.), the Giovanni Armenise-Harvard Foundation’s HMS Junior Faculty Grants Program, the Mckenzie Family Charitable Trust Systems Biology Fellowship Fund, the William F. Milton Fund, the Harriet Sugar Toibin Charitable Gift Annuity – Gene Intervention Research, NSF grants 1715184 and IOS-1452557, NIH grants R21 HD072481, R01 GM122928 and U01 GM103804 (to A.H.D.), and the Novartis fellowship (to J.E.). We thank Jeehae Park for her contribution of the fixed y[1] w[67c23]; attP2{hbP2-LacZ} embryos used in our analyses.

