## Supplementary material for "A mutation in the *Drosophila melanogaster eve* stripe 2 minimal enhancer is buffered by flanking sequences": Compiled Supplementary Material

**Figure S1: Binding site plots showing predicted Gt binding sites in different enhancers.** Three PWMs were used with the PATSER program to predict Gt binding sites within the enhancer sequences using a p-value of 0.001. All binding site plots have been aligned to the left. The light blue binding site present in the minWT is the gt-2 binding site deleted in the minΔgt enhancer. The binding site with overlapping bars at the right end of the extWT and extΔgt enhancers represents the Gt site predicted by the three PWMs in the flanking sequence of the extended enhancer. This binding site has been mutated in the extΔgt,Δgt enhancer.

**Figure S2: Complete expression level data for Figures 1-3.** (A) The peak *lacZ* level ratios presented throughout the study have been plotted together. A ratio of 1 has been indicated with a dashed line. The p-values resulting from t-test comparisons between different peak *lacZ* level ratios, corrected with the Bonferroni test (p-value from each t-test multiplied by total number of comparisons in the plot), have been included on top of the data points. (B) The same ratios as in A were plotted with each stain labelled with a unique symbol. The average ratio for all stains has been included as a red diamond. (C) The number of embryos from each transgenic line included in the plots with the peak *lacZ* level ratios has been sorted per stain.

**Figure S3: Proof-of-principle of *eve* stripe 2 expression normalization by *eve* stripe 1**

(A) Peak *eve* stripe 1 intensity and peak *eve* stripe 2 intensity follow a linear relationship, indicating that stripe 1 is an appropriate measure for normalization. Three different stains of the WT *eve* locus were analyzed (stain 1: n=10, stain 2: n=6 and stain 3: n=5). The r-squared coefficient of the linear relationship is 0.9220. The genotypes were as follows: stain 1: *y*[1] *w*[67c23]; attP2{hbP2-LacZ}, stain 2: *y*[1] *w*[67c23] ; attP{*eve*-TER94 construct} (Fujioka *et al.* 2013), stain 3: *y*[1] *w*[67c23]; attP2{evep-eve37}. (B) Normalized peak stripe expression is compared across three stains of the WT *eve* locus. The expression levels do not differ significantly within stripes across multiple stains. Filled circles represent the mean expression level within a stain and open circles are individual embryo peak expression in stain 1: n=10, stain 2: n=6 and stain 3: n=5.

**Figure S4: *Eve* locus expression data for Figure 4**. The left panels show the average stripe boundaries for *eve* stripes 3-7; the error bars represent the standard error of the mean. The right panels show the *eve* peak expression level for the corresponding stripe in the WT *eve* locus and the Δgt *eve* locus. Filled circles represent the mean expression level and open circles are individual embryo peak expression. (A) Stripe 3 boundaries and peak expression levels do not change significantly in the Δgt *eve* locus compared to the WT *eve* locus (p=0.5545). (B) Stripe 4 boundaries significantly shift anteriorly at a point on the dorsal side of the embryo indicated by the asterisk (p=0.309). However, *eve* stripe 4 peak expression levels do not change significantly in the Δgt *eve* locus (p=0.263). (C) Stripe 5 boundaries do not shift significantly, but there is a significant increase in *eve* expression in the Δgt *eve* locus (p=0.0047). During this time in development, stripe 5 is the most dynamic, and although embryos were age controlled, there may be small differences in age resulting in the observed increase. (D) Stripe 6 boundaries do not shift, but there is a significant increase in *eve* expression in the Δgt *eve* locus (p=0.0181). (E) Stripe 7 boundaries and levels in Δgt *eve* locus do not change significantly compared to the WT *eve* locus.

**Figure S5: Binding site plots showing changes to predicted binding sites in the *eve* locus after CRISPR deletion of gt-2.** To identify all predicting binding sites, we used multiple available PWMs for each *eve* regulator (Bicoid, Caudal, Giant, Hunchback, Huckebein, Knirps, Krüppel, Tailless, and Zelda), totalling 87 PWMs. These PWMs were used with the PATSER program to predict binding sites within the *eve* minimal stripe 2 enhancer sequences using a p-value of 0.001. All binding site plots have been aligned to the left. Note that removal of gt-2 in the Δgt *eve* locus could have potentially disrupted predicted tailless, caudal and huckebein sites. We did notice the addition of a SNP on the 3' end of the Δgt eve locus, which may create an additional Caudal binding site (File S1).

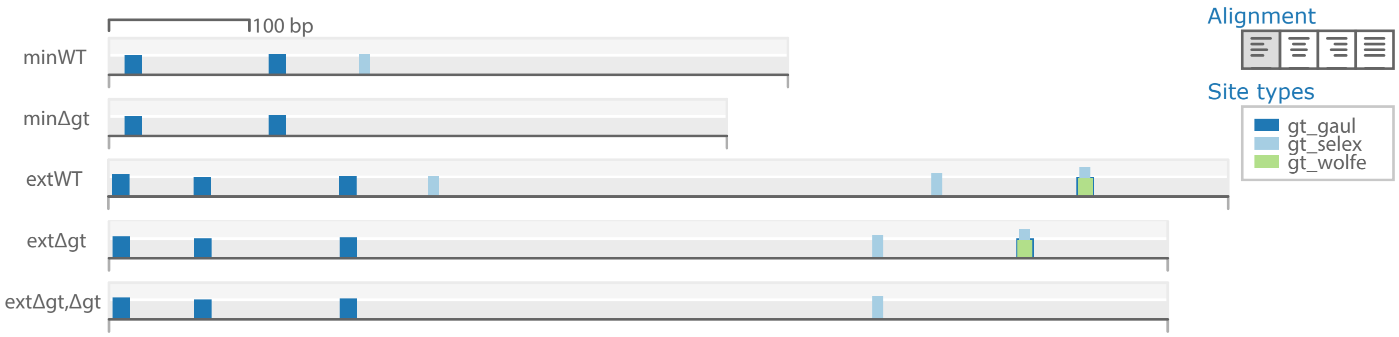
100 bp
minWT
minΔgt
extWT
extΔgt
extΔgt,Δgt
Alignment
Site types
gt_gaul
gt_selex
gt_wolfe

A

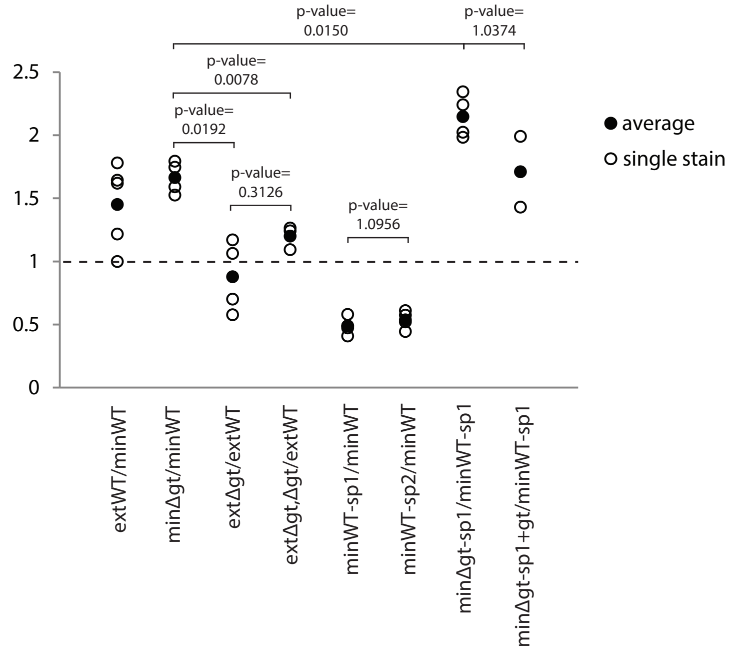


B

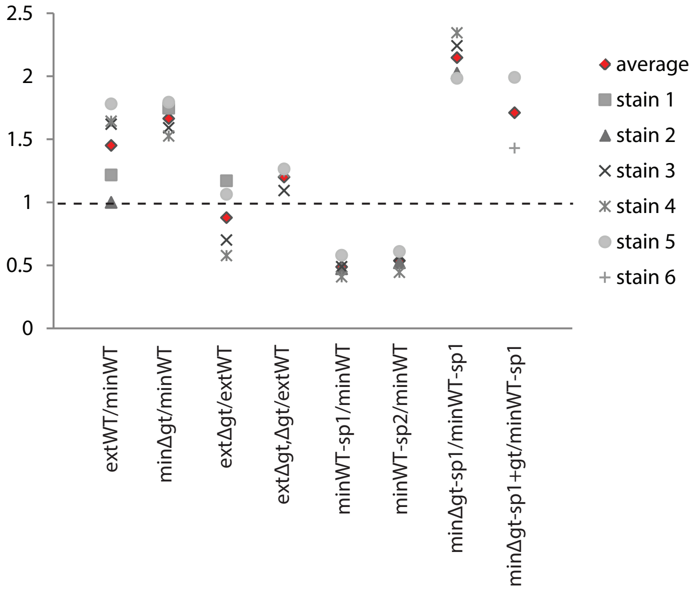


C

| **Transgenic line** | **Stain 1** | **Stain 2** | **Stain 3** | **Stain 4** | **Stain 5** | **Stain 6** |
|---|---|---|---|---|---|---|
| minWT | 9 | 15 | 13 | 13 | 33 | |
| extWT | 10 | 12 | 21 | 19 | 24 | |
| minΔgt | 10 | | 9 | 36 | 23 | |
| extΔgt | 14 | | 11 | 8 | 12 | |
| minWT-sp1 | | 9 | 11 | 21 | 23 | 11 |
| minWT-sp2 | | 13 | 14 | 11 | 21 | |
| minΔgt-sp1 | | 14 | 24 | 9 | 29 | |
| extΔgt,Δgt | | | 22 | 12 | 19 | |
| minΔgt-sp1+gt | | | | | 25 | 11 |

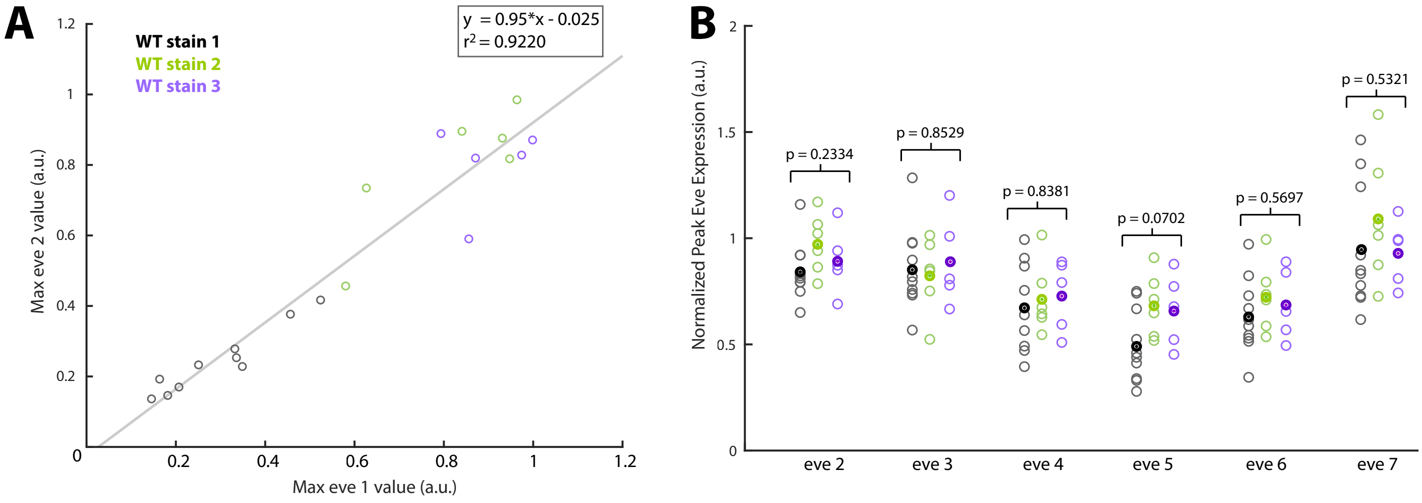
A
WT stain 1
WT stain 2
WT stain 3
y = 0.95*x - 0.025
r2 = 0.9220
Max eve 2 value (a.u.)
Max eve 1 value (a.u.)
B
Normalized Peak Eve Expression (a.u.)
p = 0.2334
p = 0.8529
p = 0.8381
p = 0.0702
p = 0.5697
p = 0.5321
eve 2
eve 3
eve 4
eve 5
eve 6
eve 7

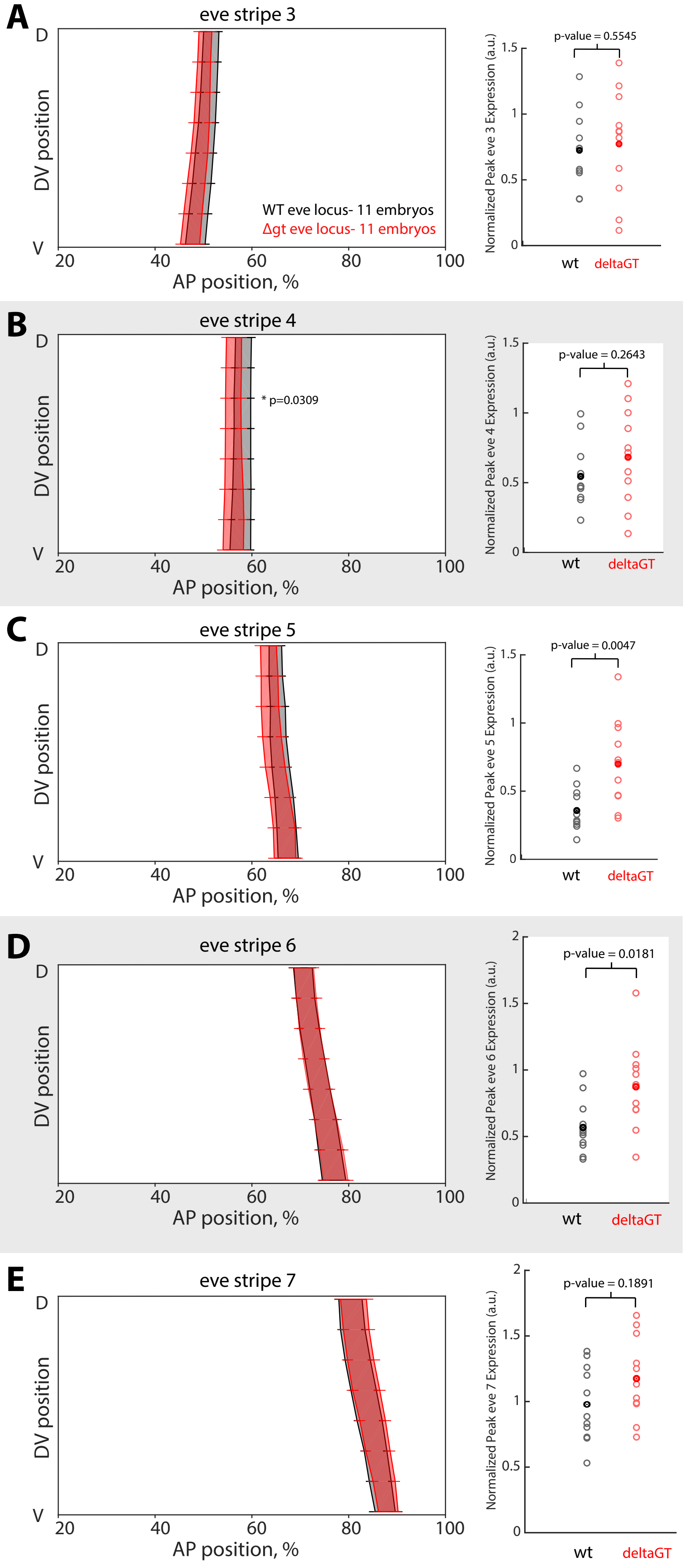
A
eve stripe 3
D
V
DV position
WT eve locus- 11 embryos
Δgt eve locus- 11 embryos
20 40 60 80 100
AP position, %
p-value = 0.5545
Normalized Peak eve 3 Expression (a.u.)
wt deltaGT
B
eve stripe 4
* p=0.0309
p-value = 0.2643
Normalized Peak eve 4 Expression (a.u.)
C
eve stripe 5
p-value = 0.0047
Normalized Peak eve 5 Expression (a.u.)
D
eve stripe 6
p-value = 0.0181
Normalized Peak eve 6 Expression (a.u.)
E
eve stripe 7
p-value = 0.1891
Normalized Peak eve 7 Expression (a.u.)

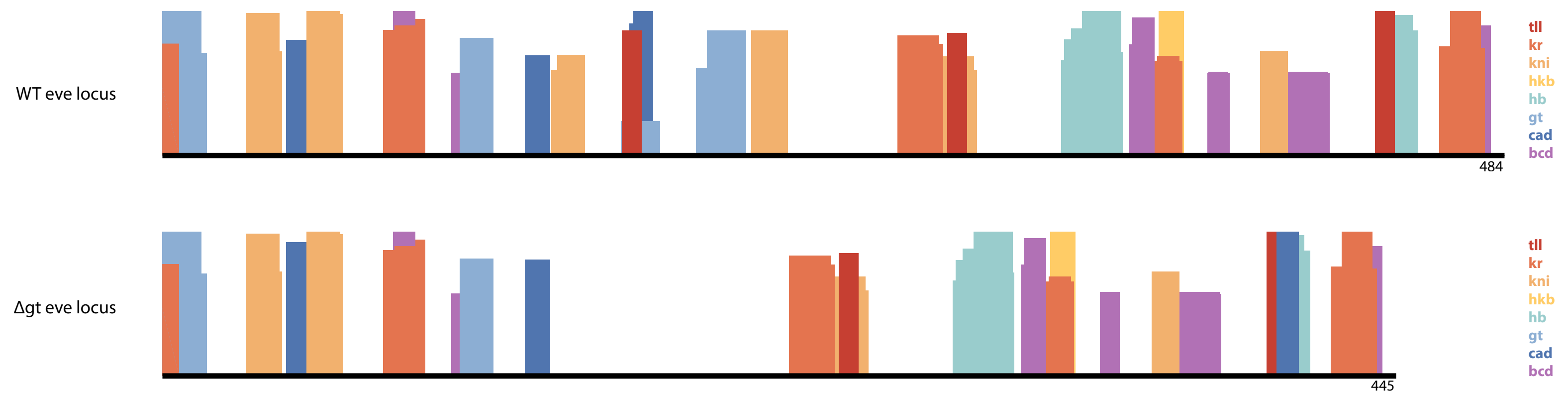
WT eve locus
484
Δgt eve locus
445
tll
kr
kni
hkb
hb
gt
cad
bcd
tll
kr
kni
hkb
hb
gt
cad
bcd
